# Convergence of socio-ecological dynamics in disparate ecological systems under strong coupling to human social systems

**DOI:** 10.1101/296202

**Authors:** Ram Sigdel, Madhur Anand, Chris T. Bauch

## Abstract

It is widely recognized that coupled socio-ecological dynamics can be qualitatively different from the dynamics of social or ecological systems in isolation from one another. The influence of the type of ecological dynamics on the dynamics of the larger socio-ecological system is less well studied, however. Here, we carry out such a comparison using a mathematical model of a common pool resource problem. A population must make decisions about harvesting a renewable resource. Individuals may either be cooperators, who harvest at a sustainable level, or defectors, who over-harvest. Cooperators punish defectors through social ostracism. Individuals can switch strategies according the costs and benefits of harvesting and the strength of social ostracism. These mechanisms are represented by a differential equation for social dynamics which is coupled to three different types of resource dynamics: logistic growth, constant inflow, and threshold growth. We find that when human influence is sufficiently weak, the form of natural dynamics leaves a strong imprint on the socio-ecological dynamics, and human social dynamics are qualitatively very different from natural dynamics. However, stronger human influence introduces a broad intermediate parameter regime where dynamical patterns converge to a common type: the three types of ecological systems exhibit similar dynamics, but also, social and ecological dynamics strongly mirror one another. This is a consequence of stronger coupling and is reminiscent of synchrony from other fields, such as the classic problem of coupled oscillators in physics. Socio-ecological convergence has implications for how we understand and manage complex socio-ecological systems. In an era of growing human influence on ecological systems, further empirical and theoretical work is required to determine whether socio-ecological convergence is present in real systems.

## 1. Introduction

Common pool resources (CPR) are natural or human-made resources available to everyone for consumption, but where excluding individuals is difficult and where one person’s use subtracts from another’s use, leading to a tendency for the resource to be over-exploited (Ostrom, 2007, 2009). Some examples of common pool resources are certain fisheries, forests, groundwater basins, pastures, and irrigation systems. Until the 1980s, a commonly held assumption was that users of common pool resources will overexploit the resource in the absence of state control or full privatization (Cox et al., 2010). However, Elinor Ostrom showed that a third approach was possible, and was the first to formulate eight design principles to regulate the common pool resources through self-organization processes in which the government has only an indirect role, and emergent phenomena including social norms and graduated sanctions help protect the resource (Ostrom, 2015). Subsequent research has refined and confirmed the insight that CPR users can develop self-governing institutions through communication among themselves and stop the commons from being over-harvested, with little or no dependence on the state (Cox et al., 2010; Sarker et al., 2015).

Forests are often used as a case study in CPR. On the global scale, forest cover has been in overall decline over the past few centuries as forests are cleared for their timber and to make way for agriculture. However, past decades have seen the reversal of this trend in some countries, in a ‘forest transition’ to net re-forestation (Lambin and Meyfroidt, 2011; Pagnutti et al., 2013). This has been driven both by improving crop yields, but also in some cases by socio-ecological interactions, which slow down deforestation and stabilize forest cover (Lambin and Meyfroidt, 2010). In several developing countries, local ownership of forests came into effect after a government policy of decentralization. Some community based forestry approaches include: the Community Forestry, Nepal; Forest Councils, Kumaon; and the Joint Forest Management in India. These programs are intended to support sustainable resource management by maintaining the local population’s right to participation in decision-making, in contrast to other approaches such as the Parks and Peoples program in Nepal that do not guarantee the local population’s rights and participation (Agrawal and Gupta, 2005; Agrawal and Ostrom, 2001). In Nepal, the concept of community forestry has been launched with the ambition of sustainable forest development, including poverty elimination through the reduction of various sources of social inequality. However, we note that whether community forestry reduces poverty remains an open question (Mahanty et al., 2006).

Similarly, over-fishing exhibits features of the CPR problem. In many commercial fishing methods that catch multiple species, by-catch of threatened species is a major source of depletion of many fish species. Also, commercially important fish species are themselves under threat due to overfishing. Historical examples from fisheries verify the frequent failures of open access regimes and the successes of the community-based management systems (Uchida and Wilen, 2004; Berkes, 2005).

Standard competition theory assumes individuals compete to exploit the common pool resource and thus reduce resource levels to their lowest possible level (Tilman, 1994). Theoretical models of CPRs have extended classical work to include social norms, ostracism and sanctions that favour cooperation (Lade et al., 2013; Iwasa and Lee, 2013; Tavoni et al., 2012; Sigdel et al., 2017). Individuals who cooperate with one another may choose to punish defectors in the group who attempt to over-exploit the resource, thus ensuring the resource is sustainably harvested. From a socio-ecological perspective, CPR problems exemplify a coupled socio-ecological system (or human-environment system), where the dynamics of a social system influence an ecological systems, and *vice versa* (Innes et al., 2013; Henderson et al., 2013). Previous research has explored how the tension between social norms that tend to drive population opinion to conformity, and conservation priorities that become stronger as a resource becomes more rare, compete with one another to generate complex dynamics (Sigdel et al., 2017). Other research explores how changing the time horizons and discounting rates that individuals use in conservation decisions can have differing impacts on resource level and stability in socio-ecological forest models (Henderson et al., 2016), and compares how different policies can result in different socio-ecological dynamics, some of which are more productive than others (Henderson et al., 2013). The role of population heterogeneity in the efficiency of graduated sanctions has been explored in mathematical models (Iwasa and Lee, 2013), while agent-based models enable researchers to incorporate greater socioeconomic and demographic complexity to explore human-wildlife interactions (An et al., 2005; Marley et al., 2017).

In coupled socio-ecological systems, regime shifts may be present and are often analyzed through mathematical models. Human feedback has a crucial role in shaping socio-ecological regime shifts. Feedback of declining natural dynamics on human opinion and behaviour may not only prevent the system from collapsing through a critical transition, but also creates several options through which the resources can survive (Bauch et al., 2016). Regime shifts in CPR can be stopped under cooperative harvesting practices where resource over-exploitation due to non-cooperative harvesters is controlled through social ostracism. Such regime shifts are not displayed in ecological subsystems in isolation from human dynamics (Lade et al., 2013; Sigdel et al., 2017). Surprisingly, increasing the resource flow sometimes results in a regime shift from a high level of cooperation to violation of socials norms and collapse of the resource (Lade et al., 2013). In this case, as the defectors are accruing more benefit from a larger resource flow, it favors the cooperators to switch their strategy (Tavoni et al., 2012). Other research explores how gradual change in an external driver can result in abrupt change in the threshold system, if ecosystem dynamics are defined by a threshold in which a state and the ecosystem’s driver are not linearly co-related (Ratajczak et al., 2014).

Forests, fisheries, groundwater basins, and other CPR problems are represented by diverse natural dynamics. Some systems may be open to restocking or recolonization from outside while others are relatively closed. Other systems may exhibit threshold behaviour where the resource abundance must exceed a certain threshold in order to remain viable. Although CPR problems have been relatively well-studied in the mathematical modelling literature, comparative analyses of how different types of resource dynamics influence socio-ecological dynamics are infrequent. Our objective in this paper is to study how the type of resource dynamics influences outcomes in a model of a socio-ecological system. We develop three ordinary differential equation models of common-pool resource dynamics, corresponding to three different types of natural dynamics. Individuals either cooperate to harvest sustainably (and punish those who do not through social ostracism) or they defect and over-exploit the resource. We explore and compare model dynamics through stability analysis and numerical analysis. The model is described in the next section.

### 2. Model Development

### 2.1. Model 1

Consider a resource limited by a carrying capacity, where *F* ∈ [0, 1] is the current level of the resource compared to its maximal carrying capacity *F* = 1. Let *R*, *µ* and *n* be the growth rate, natural death rate and the number of resource users per year, respectively. We consider a harvesting community in which individuals are divided into two opinion groups: *x* is the proportion of cooperators and 1 − *x* is the proportion of defectors. We denote *e_c_* and *e_d_ > e_c_* as the harvesting effort for cooperators and defectors respectively. We assume logistic growth of resource so that the new resource is created based on the resource level, such as through population reproduction. We assume that the resource is removed in two ways: natural death and harvesting. Thus, the rate of change in resource dynamics can be written as:

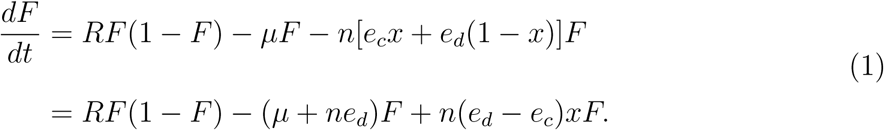

In the above equation, the last term *n*[*e_c_x* + *e_d_*(1 − *x*)] represents total harvesting intensity (Lade et al., 2013).

Using a similar approach to Sigdel et al. (2017), an individual adopts one of the two opinions, either to cooperation (ℂ) or defection (ⅅ). Based on the two strategies, individuals get sample one another at the rate *κ*. In this case, an individual adopting strategy ℂ samples other individuals adopting either of the strategies ℂ or ⅅ. Let *π*(ℂ) be the payoff function of cooperators and *π*(ⅅ) is of defectors.

Before switching their strategies, an individual in either group compares the payoff gain or loss received by adopting the same strategy, and may switch strategies if the payoff for switching is attractive enough. If *π*(ℂ) *> π*(ⅅ) then the one playing ⅅ switches to strategy ℂ with probability *pU_C_*, where *U_C_* = *π*(ℂ) − *π*(ⅅ) *>* 0 is the net gain in payoff by switching to strategy ℂ and *p* is a proportionality constant (we require a sufficiently small timestep such that the probability is always less than 1). Therefore, (1 − *x*) defectors at any given time become cooperators at the rate (1 − *x*)*κxpU_C_*. Similarly if *U_D_* = *π*(ⅅ) − *π*(ℂ) *>* 0 then *x* cooperators becomes defectors at the rate *xκ*(1 − *x*)*νpU_D_* where, *U_D_* is the net gain in payoff by adopting strategy ⅅ than adopting strategy ℂ and *ν* is the scaling constant. For convenience, we absorb *ν* into *κ*. Combining the above two transition terms gives:

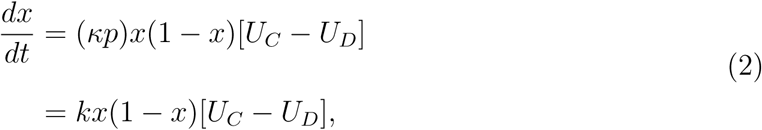

where *k* = *pk*′.

The payoff function depends on the net benefits of harvesting, which is the payoff from resource production through harvesting minus the cost of harvesting per unit effort. In addition, defectors suffer reduced payoff due to social ostracism according to an ostracism function. In this case, we assume the impact of ostracism on the defector payoff depends linearly on the number of cooperators, *x*. In particular, the ostracism function takes the form 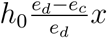, where *h*_0_ is a proportionality constant controlling the overall magnitude of ostracism and the ratio 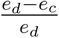 determines the strength of the ostracism as it relates to harvesting efforts. Hence, the payoff function for two groups can be written as:

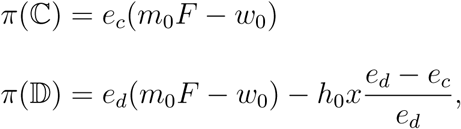

where *m*_0_ and *w*_0_ are the average resource productivity rate and the harvesting cost, respectively.

Substituting these values into equation (2) and replacing 2*m*_0_, 2*w*_0_, 2*h*_0_ by *m*, *w* and *h* respectively gives us the following equation to represent strategy dynamics:

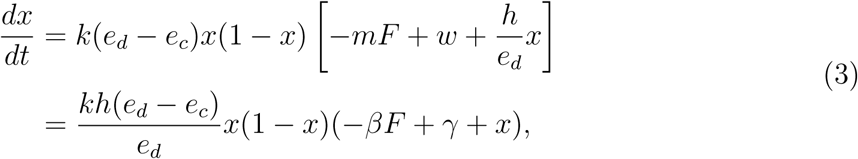

where 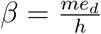 and 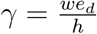.

Let us non-dimensionalize the system given by equations (1) and (3). As the dependent variables *x* and *F* are fraction of one, they are already dimension less, thus we are left only to non-dimensionalize the independent variable *t*. Let 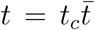 with 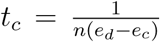 and 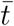 is non-dimensionalized time. Hence

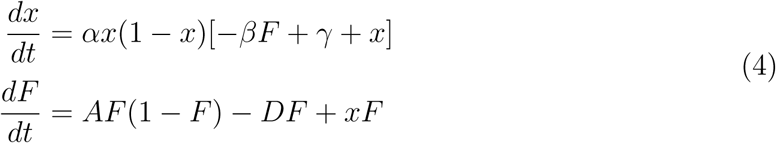

where 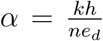, 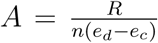 and 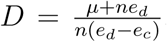. We observe that the non-dimensionalized resource dynamics depend on three major mechanisms: natural recruitment (first term involving *A*), removal due to natural death plus defectors (second term involving *D*), and increase due to the cooperators into the population (third term). These features will be shared by the next two models we develop.

### 2.2. Model 2

Instead of assuming a logistic resource growth rate as in Model 1, Model 2 considers a constant resource supply rate *λ* so that resource dynamics are given by:

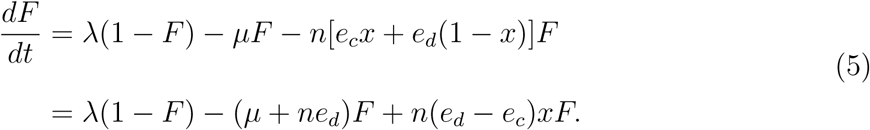

Non-dimensionalizing the resource dynamics in (5) in the same way as before gives us the social-ecological system:

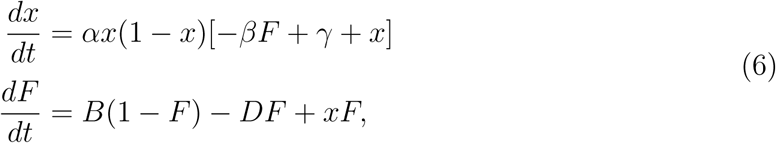

where 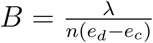.

### 2.3. Model 3

For this model, the natural resources dynamics are logistic but the recruitment constant *R* is replaced by a density-dependent recruitment rate *R*(*F*) to represent systems where growth is only possible when the resource level exceeds a fixed threshold (SI Appendix, Figure S.1). In this case the resource dynamics are given by

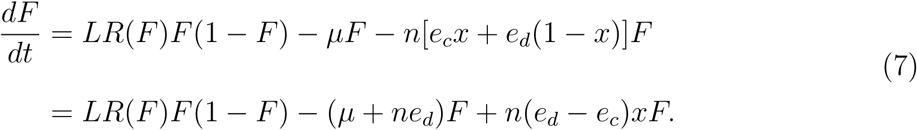

where 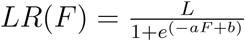 gives the resource recruitment rate in which *a* is the sharpness of the transition and *b* is the minimum transition controller. Non-dimensionalizing the resource dynamics in (7) as before yields the social-ecological system

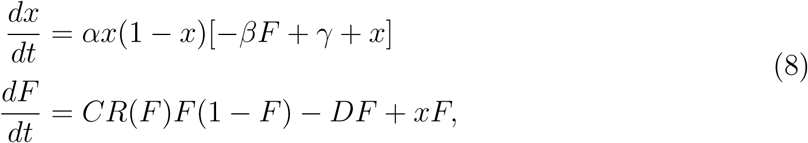

where 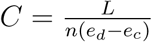.

## 3. Results

In this section we first characterize the dynamics of the three ecological subsystems in isolation from the social subsystem, before showing how these dynamics change under the addition of weak or strong social feedback.

### 3.1. Ecological system in isolation

From special cases of the coupled systems of equations in (4), (6) and (8), we first analyze the following equations for resource dynamics in isolation from social dynamics:

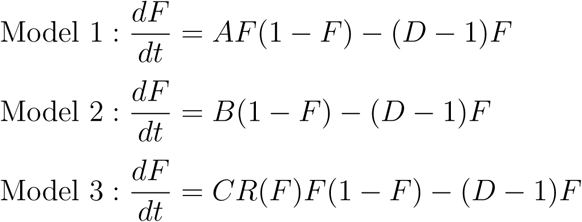

### 3.1.1. Steady states and their stability

Model 1 has two steady states, *F*^∗^ = 0 and 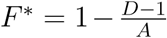. The first is stable when *A < D* − 1 and unstable otherwise, while the second is stable when *A > D* − 1 and unstable otherwise.

Note that the second steady state is stable when it is biologically meaningful. Model 2 has only one steady state 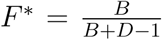 that is stable when it is biologically meaningful (when *B* + *D >* 1). Model 3 has two steady states *F*^∗^ = 0 and *F*^∗^ = *F*_1_, where *F*_1_ is given by the solution of

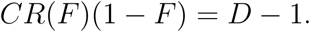

Note that *F* = *F*_1_ is biologically meaningful if *CR*(*F*_1_) *> D* − 1. Here *F* ^∗^ = 0 is stable when *CR*(0) *< D* − 1. The eigenvalue of the second steady state is zero, thus we are unable to say anything about it’s stability with only linear stability analysis.

The models yield different outputs as a function of the natural growth rate parameters *A*, *B* and *C* (Figure 1, see Table 1 for parameter values). In Model 1, there is a regime shift from the resource-free state to a stable nontrivial resource level as *A* increases. However in Model 2, only the nontrivial resource state is stable, for all values of *B*. In Model 3, when *C* is small, only the resource free state exists. However as *C* increases, a fold bifurcation introduces a critical transition beyond which both the resource-free and the nontrivial resource steady states are stable. The system converges to one state or the other depending on initial conditions.

**Table 1:**
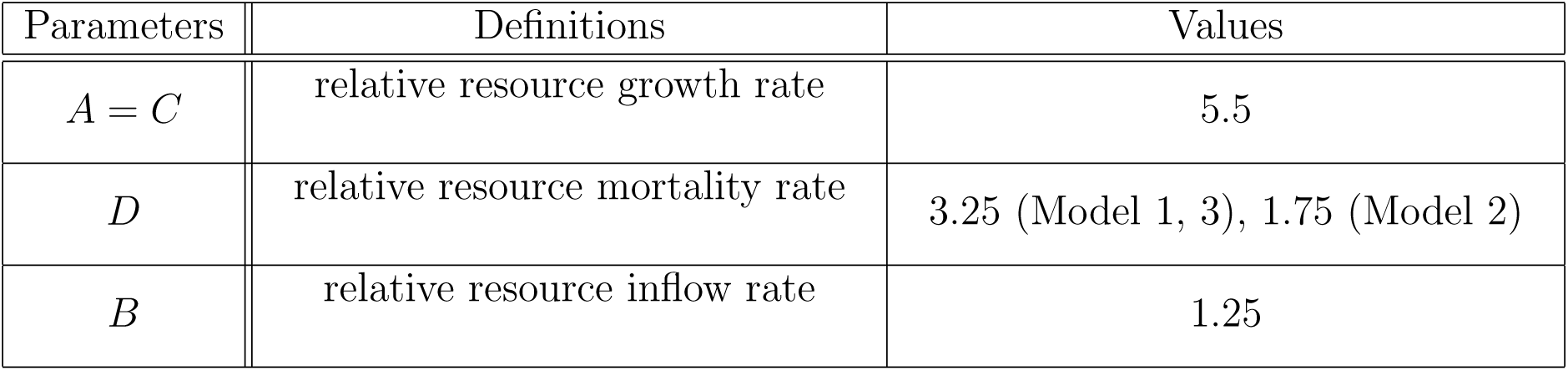
Parameters, definitions and their baseline values. These parameter values were obtained as a special case of the baseline parameter values for the coupled socio-ecological models (Table 2).

| Parameters | Definitions | Values |
| --- | --- | --- |
| $A = C$ | relative resource growth rate | 5.5 |
| $D$ | relative resource mortality rate | 3.25 (Model 1, 3), 1.75 (Model 2) |
| $B$ | relative resource inflow rate | 1.25 |

**Figure 1:**
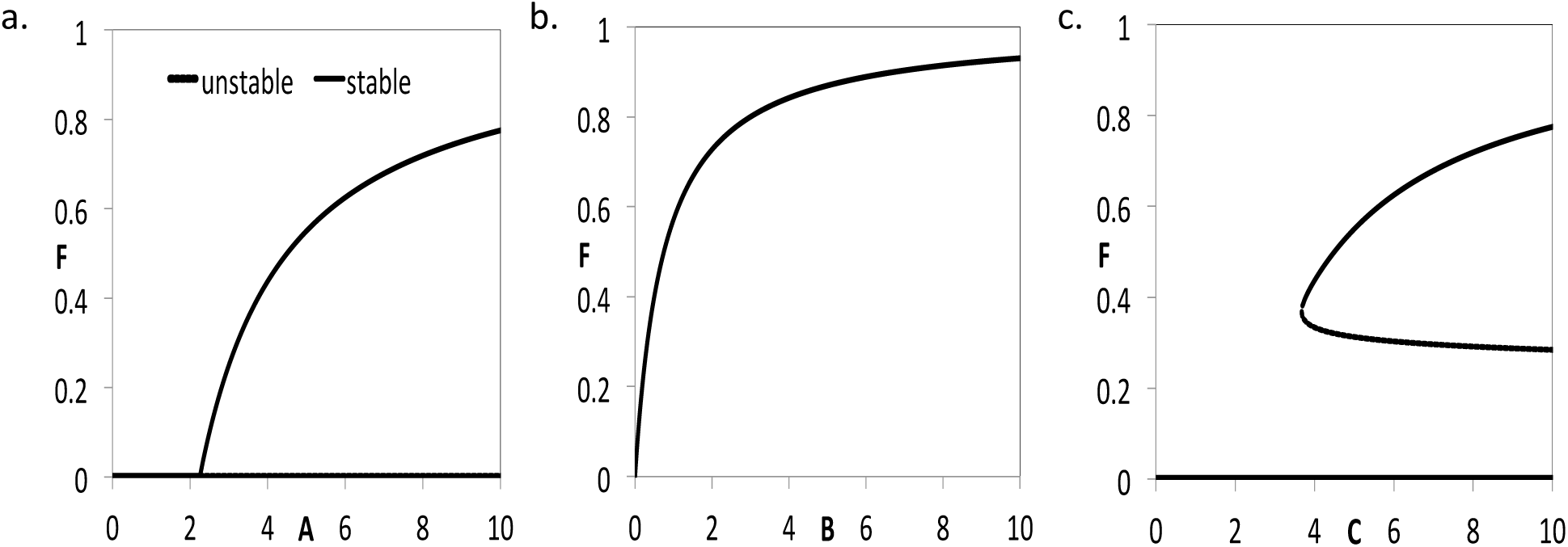
Resource growth is generally proportional to the resource growth rate parameter, with some exceptions. Stable and unstable steady states versus (a) *A*, Model 1, (b) *B*, Model 2, (c) *C*, Model 3. All other parameter are in Table 1.

As the removal rate *D* is increased, populations are reduced or collapse completely in all three models (Figure 2). In Model 1 this occurs through a transcritical bifurcation at zero resource level and in Model 3 it occurs through a fold bifurcation, corresponding to a sudden regime shift from nonzero resources to resource extinction. In Model 2, however, resources levels decline with increasing values of *D* but are never extinguished as in the other two models.

**Figure 2:**
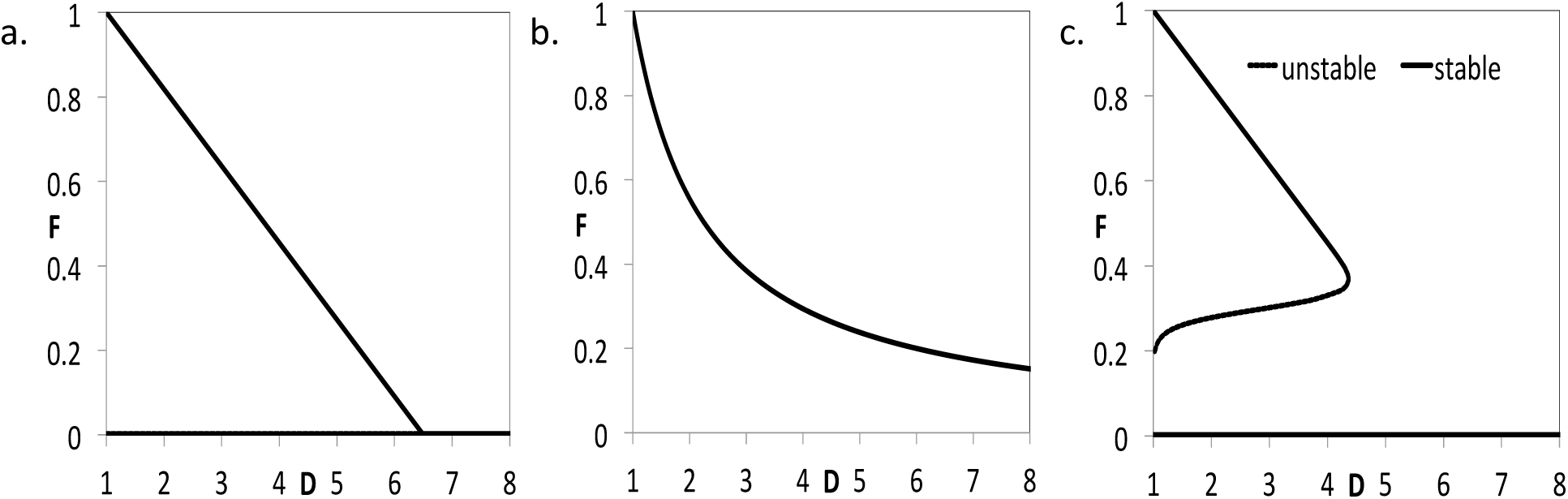
Resources decline and sometimes collapse with increasing removal rates in all three models. Steady states versus *D* for (a) Model 1, (b) Model 2, (c) Model 3. All other parameters are in Table 1.

### 3.2. Coupled socio-ecological system

In this section, we discuss the results from the coupled systems and compare it to the isolated systems for all three models with differing natural resource dynamics. We begin with discussing parameter values where human influence is relatively weak, and then we explore parameter regimes with stronger human influence.

### 3.2.1. Steady states and their stability

In the following we will let *F*_1_, *F*_2_ and *F_I_* denote the resource level in a state of all-defectors, all-cooperators and the interior state for all three models, respectively. We also let *x_I_* denote the proportion of cooperators in the interior state. For the details of local stability analysis, see SI appendix A.

#### Model 1

For system of equations (4), there are two steady states with only defectors. One of them has no resource cover

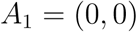

and the second one has some fraction of resource cover

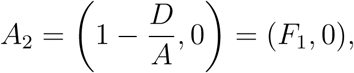

The first one is unstable and the second one is locally asymptotically stable (LAS) if the ratio of cost to resource productivity (price performance ratio) is dominated by forest cover, that is 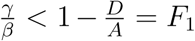. When the resource productivity is higher, the second steady state is stable. Thus the stronger factor productivity shifts the population to the all-defector state.

There are also two steady states with only cooperators. One of them has no resource cover

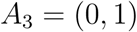

and the second one has some fraction of resource cover

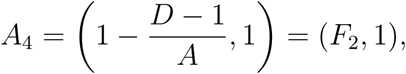

The first one is LAS if the relative fecundity rate is dominated by the relative harvesting rate of cooperators (*A < D* − 1) and the second one is LAS if 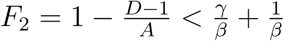.

Thus for the state without resource cover *A*_3_, a stronger harvesting rate of cooperators (*D*−1) shifts the population to the all-cooperator steady state. This means that the threshold of harvesting intensity of the cooperators is important in determining survival of the resource. The stability of the state with resource cover *A*_4_, exists for larger values of the harvesting cost (*γ*), shifting the population to the all-cooperator state.

We also have one interior steady state *A*_5_ in which both the resource cover as well as cooperators exist partially,

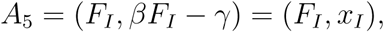

where 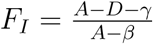, is LAS if the relative resource fecundity rate is dominated by the factor productivity and the social learning rate is dominated by the resource recruitment rate, that is *β > A* and *αx_I_*(1 − *x_I_*) *< AF_I_*, where 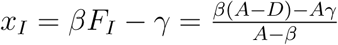 and 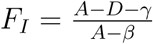.

Note that the state 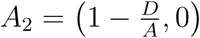 of defectors is biologically meaningful if the resource growth rate dominates the maximal harvesting rate of defectors, that is *A > D*. And the state of cooperators 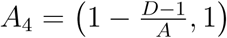 is biologically meaningful if the resource growth rate dominates the maximal harvesting rate (or the total harvesting effort) of cooperators, that is *A > D* − 1.

#### Model 2

For the system in equation (6), there is one steady state with only defector and partial resource cover

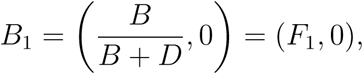

and it is LAS if 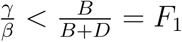. As in the first model, the larger value of resource productivity increases harvesting rate which ultimately shifts the population to the homogeneous opinion so that the state of full defectors exist.

There is one steady state with only cooperator and partial resource cover

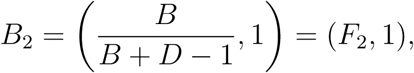

and it is LAS if 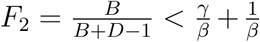. And the stability is linked with the similar constraints to the state *A*_4_ of Model 1.

We also have a steady state in which both of the opinion and the resource has intermediate values

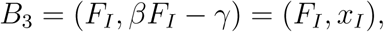

where *F_I_* is given by

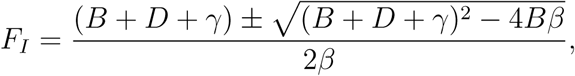

and it is LAS if *αx_I_*(1 − *x_I_*) + *x_I_ < B* + *D* and 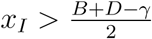.

#### Model 3

For the system in equation (8), there are two steady states with only defectors. One of them has no resource

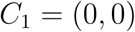

and the second one has some fraction of resource

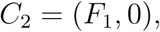

where *F*_1_ is given by

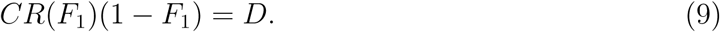

Here, the first one is unstable and the second one is LAS if the resource cover given by the equation (9) satisfy 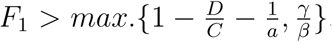. If the second constraints hold then the first and the third models are related by the same constraints of stability.

There are also two steady states with only cooperators. One of them has no resource

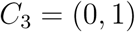

and the second one has some fraction of resource

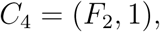

where *F*_2_ is given by

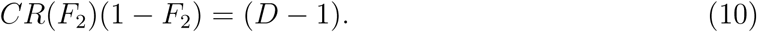

Here, the first one is LAS if the resource growth rate at the initial stage of resource cover is dominated by the harvesting intensity of the cooperators, that is *CR*(0) *< D* − 1 and the second one is LAS if the resource cover given by the equation (10) satisfy 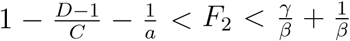. If the second constraint holds then the first and the third models are related by the same constraints of stability.

In the state of cooperators without resource cover *C*_3_, the higher harvesting intensity of cooperators shifts the population to the homogeneous opinion so that the state where the entire population of cooperators exists without existence of the resource.

We also have one interior steady state

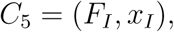

where *F_I_*, *x_I_* are given by

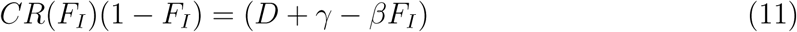

and *βF_I_* − *γ* respectively and this state is LAS, if the following conditions are satisfied:

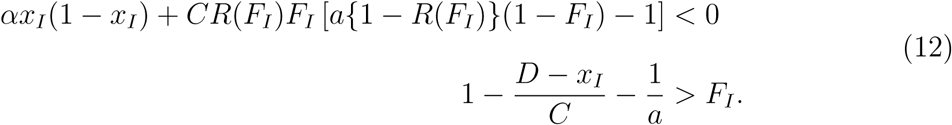

As in the first model, the state *C*_2_ of defectors is biologically meaningful if the resource growth rate dominates the maximal harvesting rate of defectors, that is *CR*(*F*_1_) *> D*. And the state of cooperators *C*_4_ is biologically meaningful if the resource growth rate dominates the maximal harvesting rate (or the total harvesting effort) of cooperators, that is *CR*(*F*_2_) *> D* − 1.

### 3.2.2. Numerical Simulations: Baseline Scenario

Baseline parameter values for the socio-ecological models (Table 2) were derived by calibrating the models to a calibration target where both *F* and *x* converge to 0.5 over a timescale of several decades (SI Appendix, Figure S.2). Our rationale was not to replicate the observed socio-ecological dynamics of particular real-world systems, but rather to create a common starting point that would make it easier to compare the dynamics of the three models under parameter variation.

**Table 2:** Variables/Parameters, their definition, ranges and baseline values.

| Variables | Definition | Ranges | – |
| --- | --- | --- | --- |
| $x$ | Proportion of cooperators | $[0, 1]$ | – |
| $F$ | Proportion of resource cover | $[0, 1]$ | – |
| Parameter | Definition | Values (Model 1, 3) | Values (Model 2) |
| $a$ | transition sharpness parameter | 50 | – |
| $b$ | transition location parameter | 15 | – |
| $\alpha$ | social learning rate | 0.8 | 0.8 |
| $\beta$ | average resource productivity rate | 7 | 7 |
| $\gamma$ | cost of harvesting | 3 | 3 |
| $A = C$ | relative resource growth rate | 5.5 | – |
| $B$ | relative resource inflow rate | – | 1.25 |
| $D$ | relative resource mortality rate | 3.25 | 1.75 |

When the resource productivity rate *β* is sufficiently small, the coupled system behaves in a similar way to the isolated system as the growth rates *A*, *B* and *C* increase (Figure 3a-c, compare to Figure 1a-c). In this case, *x* = 1 is stable across the entire parameter range. However, for a larger value of *β*, feedback from the social system splits the stable interior branches into two new branches with stable and unstable portions (Figure 3d-f).

**Figure 3:**
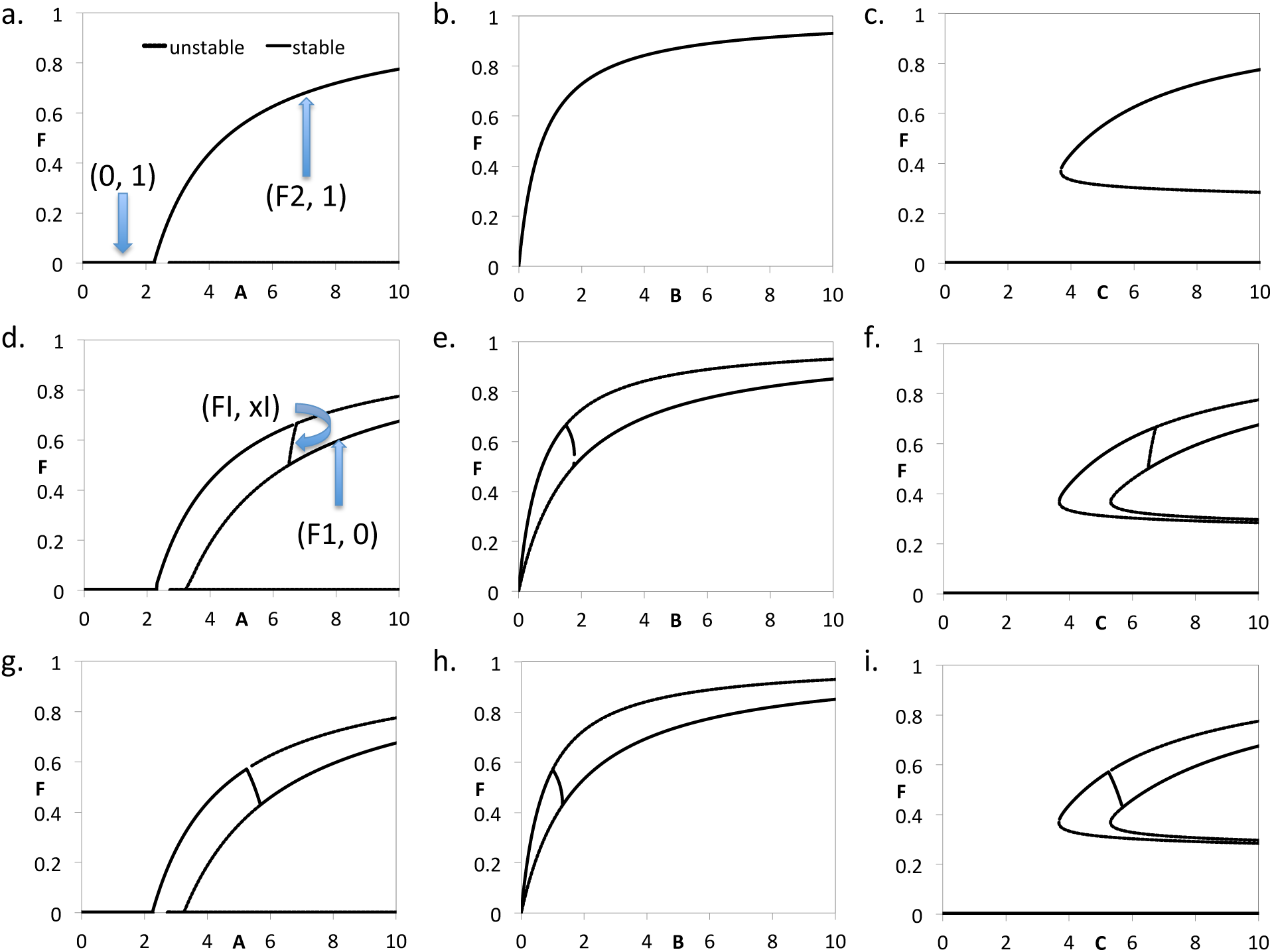
As the relative fecundity rate is increase from low to high, the stability of a state of cooperators shift to the state of the defectors. (a, d, g) Model 1; (b, e, h) Model 2; (c, f, i) Model 3. *β* = 4, a-c; *β* = 6, d-f; *β* = 7, g-i. All other parameters are in Table 2.

When the growth rate parameters are small, *x* = 1 is stable and thus the higher branch of the two new branches is stable, corresponding to more sustainable resource use. However, sufficiently high values of *A*, *B* and *C* in all three models force a switch in stability so that *x* = 0 becomes stable, along with the lower resource level branch. In models 1 and 3, the shift to the lower resource level branch is a critical transition, but not in Model 2. However, a further increase in the value of *β* causes the transition to become non-critical in all three models (Figure 3g-i). Regardless of whether the transition is critical or non-critical, the transition from the upper branch to the lower branch corresponds to a parameter regime where a growing natural growth rate causing an increase in over-exploitation and thus a (surprising) decline in the resource level.

The dependence of the three models on the removal rate *D*–which reflects both the natural death rate and the contribution to harvesting intensity from the defectors–shows similar patterns across low, medium and high values of the resource productivity parameter *β* (Figure 4). As *β* is increased, social feedback causes the interior stable branch to split into two, with the lower branch being stable for small values of *D* and the upper branch being stable for larger values of *D*. The implication is that a large enough removal rate can simulate conservationist behaviour and, across a limited range of values for *D*, result in an increase in the steady state resource level with increasing *D* instead of a decrease.

**Figure 4:**
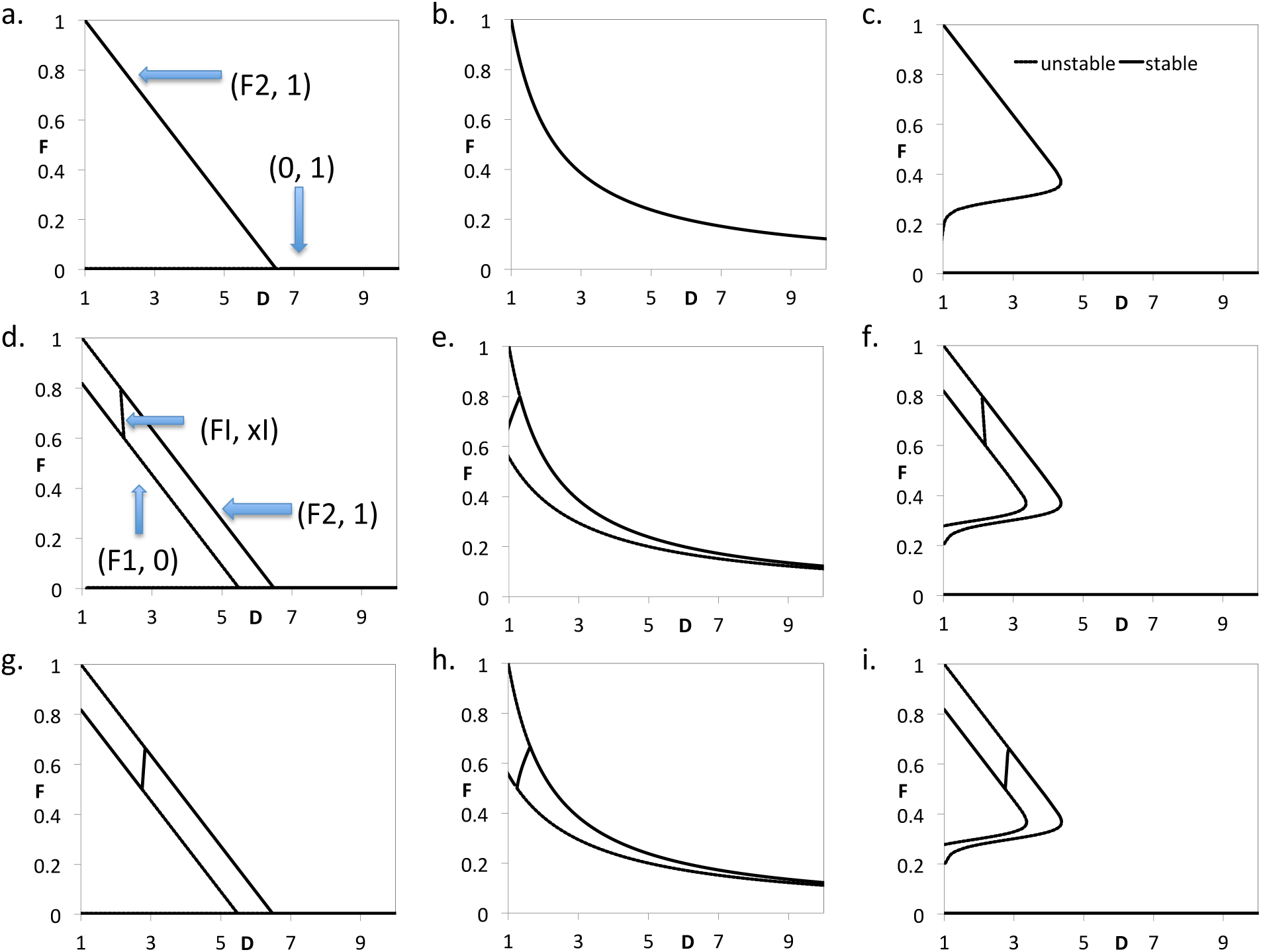
The stability shift from the state of defectors to the state of cooperators with increasing harvesting intensity of defectors but unable to protect the resource passing to collapse. (a, d, g) Model 1; (b, e, h) Model 2; (c, f, i) Model 3. *β* = 2, a-c; *β* = 5, d-f; *β* = 6, g-i. All other parameters are in Table 2.

Other outcomes are also possible in the regime of weak human influence. For instance, in models 1 and 3, if the relative resource growth rates (*A*, *C*) are too low or the relative harvesting rate of cooperators *D* − 1 is too high (*A < D* − 1, *CR*(0) *< D* − 1) then the dependence on the resource productivity rate *β* stabilizes the steady state (0, 1) of all-cooperators but no resource (SI Appendix Figure S.3).

Taken together, Figures 3 and 4 highlight the importance of the type of resource dynamics in determining outcomes in socio-ecological systems. Social opinion changes in a similar way as *A*, *B*, *C* and *D* are varied in the three models, with a tendency toward greater cooperation when the resource is threatened. However, the outcome of changing growth rates or removal rates may vary from strong resilience (Model 2) to a continuous or discontinuous transition to a state of resource extinction (Models 1 and 3, respectively). Moreover, in the isolated system, only Model 3 could exhibit a critical transition in the resource level, while in the coupled socio-ecological system, Model 1 may also exhibit a critical transition.

However, the baseline scenario corresponds to a situation where human influence is still relatively weak, since the degree of splitting of the interior stable branch as resource productivity *β* increases is relatively small. In many real systems, the presence or absence of humans can determine natural dynamics more strongly. Hence, in the next subsection we explore the scenario of strong human influence.

### 3.2.3. Numerical Simulations: Strong Human Influence

Next we explore the case of strong human influence by making natural forces relatively weaker through parameter value changes. Working from the original dimension-bearing parameterization, for models 1 and 3 we reduced the growth rate *R* and *L* from 0.022/year to 0.01/year, and we reduced the natural death rate of the resource *µ* from 0.008/year to 0.001/year. For Model 2, we reduced the resource inflow rate *λ* from 0.005/year to 0.0025/year and the natural death rate of resource *µ* from 0.002/year to 0.00025/year. These changes are reflected in the re-parameterized values of *A*, *B*, *C* and *D* and these reductions in the natural growth and death rates act to make human influence relatively stronger.

In this case, the trends observed in Figures 3 and 4 are exaggerated to the point where the resource levels begin to diverge from the general pattern observed in the isolated systems, across most of the parameter regimes tested for *A*, *B*, *C* and *D* (Figure 5). The distinct patterns of the isolated systems begin to be replaced by a broad intermediate parameter range with similar features across all three models, where the interior steady state is stable but the resource is decreasing with increasing natural growth rates (Figure 5a-c), and increasing with the increasing removal rate (Figure 5d-f). As in Figures 3 and 4, social dynamics evolve in qualitatively similar ways across these parameter ranges, with *x* transitioning between *x* = 0 and *x* = 1 as the intermediate parameter regime is traversed, but outside of this intermediate parameter regime, the type of natural dynamics can still make the difference between resilience and collapse (Figure 7).

**Figure 5:**
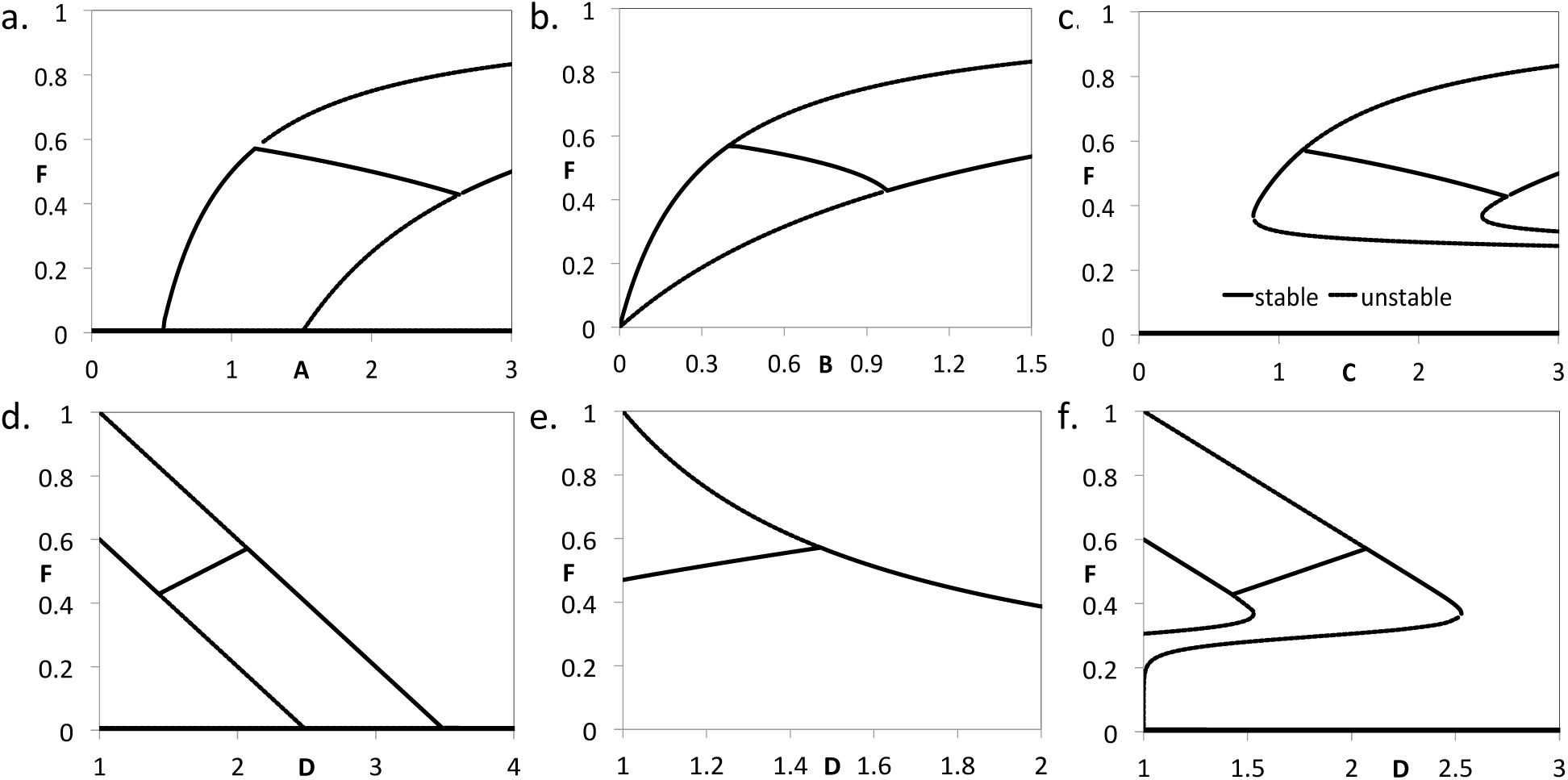
Stronger human influence relative to natural influence cause diverse effects in resource dynamics. (a, d) Model 1; (b, e) Model 2; (c, f) Model 3. All other parameters are in Table 2.

In a similar exercise, working from the reduced values of the natural parameters, let us double the effort of defectors. This changes the parameter values to: *α* = 0.4, *β* = 14, *γ* = 6, *A* = *C* = 1.1, *D* = 1.22 (Model 1, 3), *B* = 0.27, *D* = 1.14 (Model 2). Compared to Figures 5-6, the dependence of resource level on *A*, *B*, *C* and *D* is generally similar, although a few differences emerge (SI Appendix, Figure S.4). Most notably, the interior steady state is more constant with respect to changes in *A*, *B*, *C* and *D* and the intermediate range is broader.

Making social ostracism sufficiently strong can make the socio-ecological dynamics converge to the patterns observed in the isolated systems for models 1 and 3 but not for Model 2. If we double the strength of social ostracism we obtain *α* = 1.6, *β* = 3.5, *γ* = 1.5. The dependence of resource dynamics on *A*, *B*, and *C* is similar, with broad parameter regimes where increasing *A*, *B* and *C* or decrease *D* causes resource levels to vary only in a narrow range of values due to social feedback on the natural systems (Figure 7a-c). However, if the strength of social ostracism is tripled, the strength of social norms tends to maintain a stable upper branch in resource levels for a broader range of parameter values (Figure 7d-f). The *x* = 0 branch becomes unstable across the entire parameter regime in this situation, meaning there are no defectors and the population converges to an all-cooperator steady state. This tends to restore the model output so that it once again resembles the output of the isolated system in models 1 and 3 (Figure 7d, f). However, a critical transition emerges for Model 2 (Figure 7e).

Increasing the social learning rate can have striking effects on the system, leading to convergence of dynamics between the three systems across a broad parameter regime (Figure 8). It also tends to destabilize the system, leading to oscillations in the prevalence of cooperators and in the resource level. For these diagrams we increased the social learning rate from *α* = 0.8 to *α* = 5 (Model 1 and 3) and *α* = 8 (Model 2). This destabilizes the interior steady states through Hopf bifurcations, causing the emergence of stable limit cycles for all three models (Figure 8a-c). This is also observed in the corresponding bifurcation diagrams for *x* versus *A*, *B* and *C* (Figure 8d-f). In this parameter regime, human and natural dynamics mirror one another through their shared characteristic of stable limit cycles. This contrasts with different dynamics in the human and natural subsystems observed in Figures 5 and 6. The oscillations in the human system are also more extreme than the oscillations in the natural system, on account of the slower dynamics in the natural system. Time series show damped oscillations in resource level and prevalence of cooperation, and these oscillations persist for a longer period of time in Model 2 than in the other models (SI Appendix, Figure S.5).

**Figure 6:**
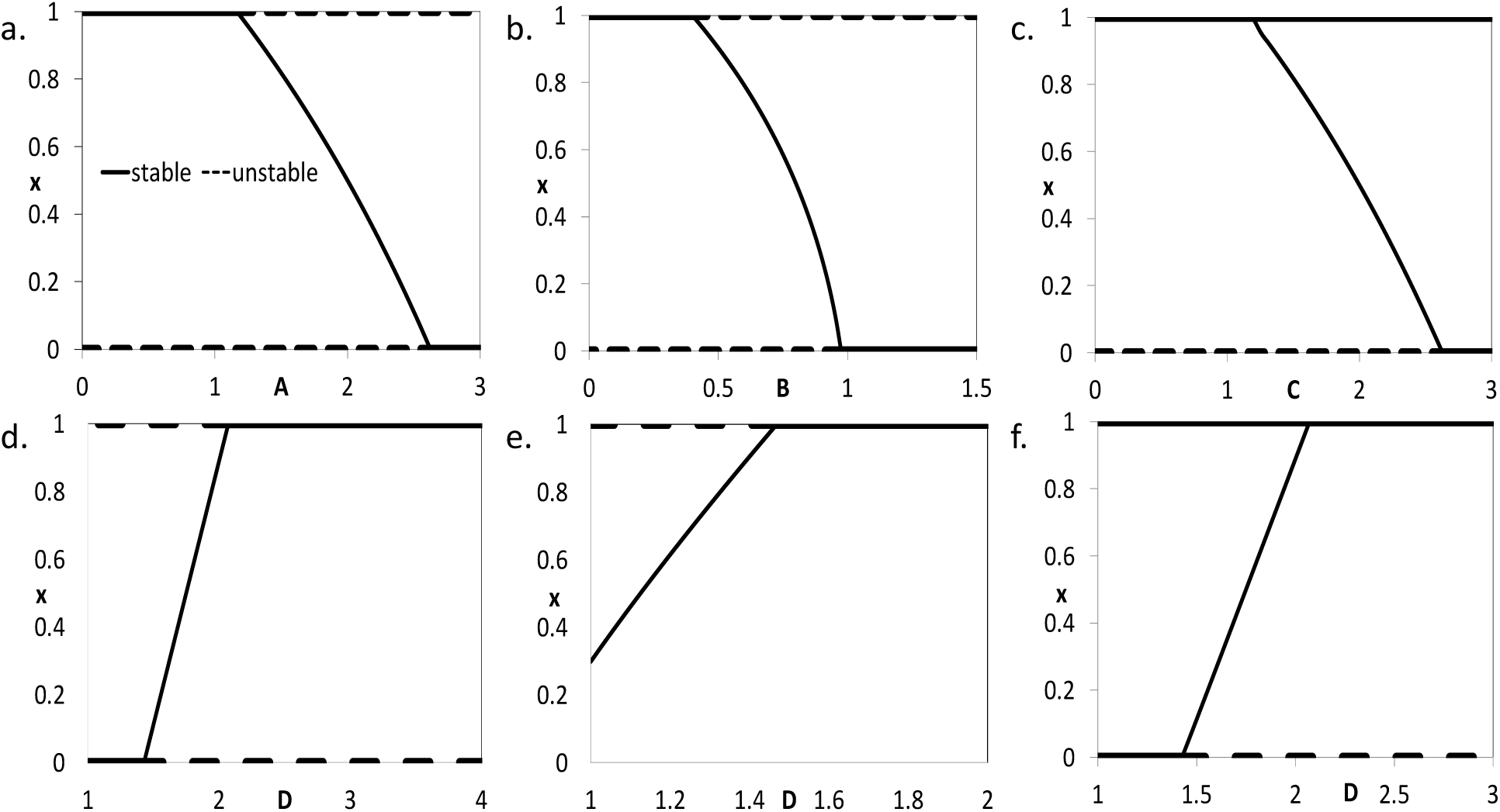
Stronger human influence relative to natural influence cause diverse effects in opinion dynamics. (a, d) Model 1; (b, e) Model 2; (c, f) Model 3. All other parameters are in Table 2.

**Figure 7:**
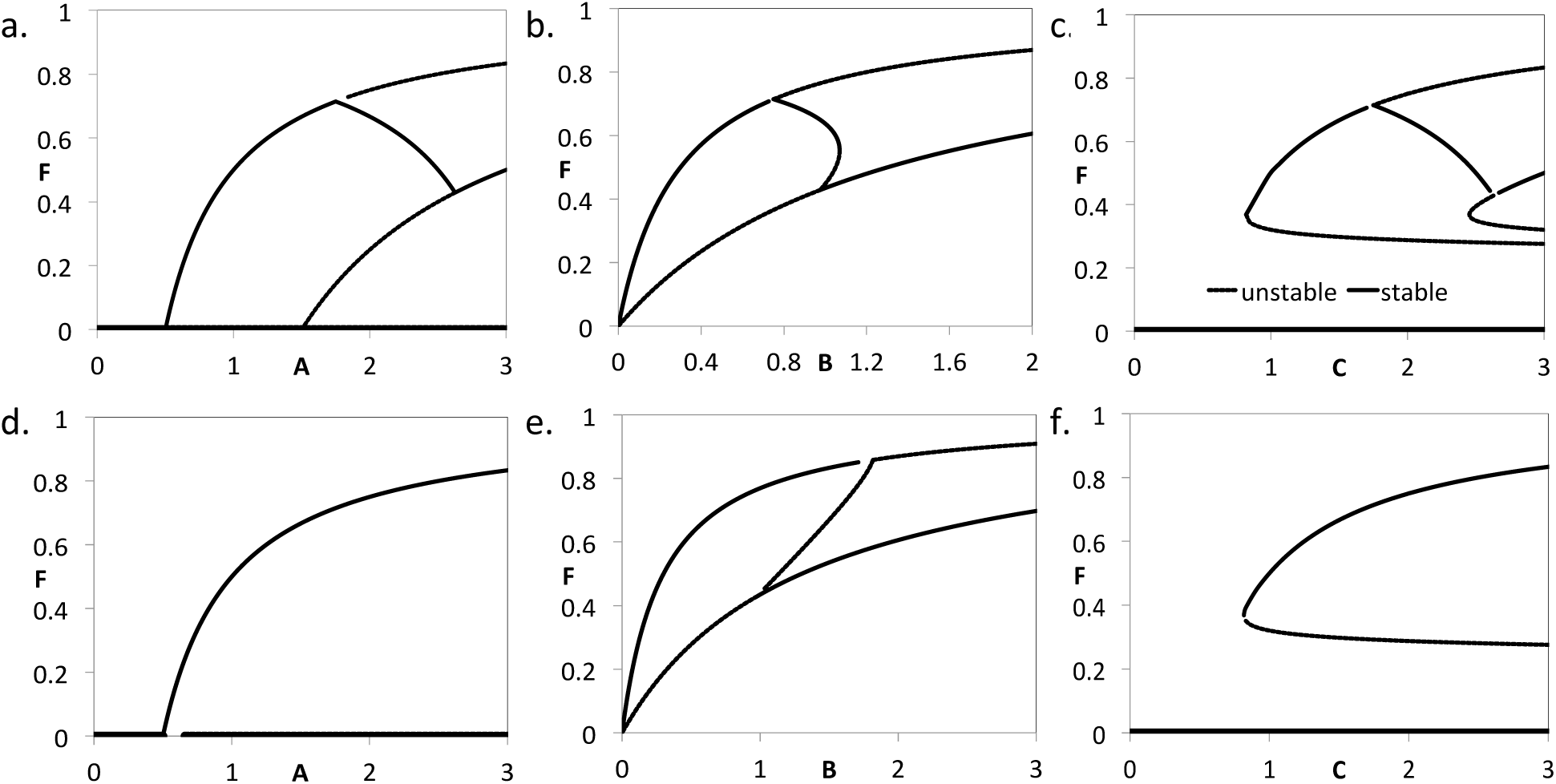
Increasing ostracism helps to stop defection and thus the population of only cooperators exist in Model 1 and 3. The figures in the first row comes from the two times larger values of the social ostracism whereas the figures in the second row from the three times larger value. (a, d) Model 1; (b, e) Model 2; (c, f) Model 3. All other parameters are in Table 2.

**Figure 8:**
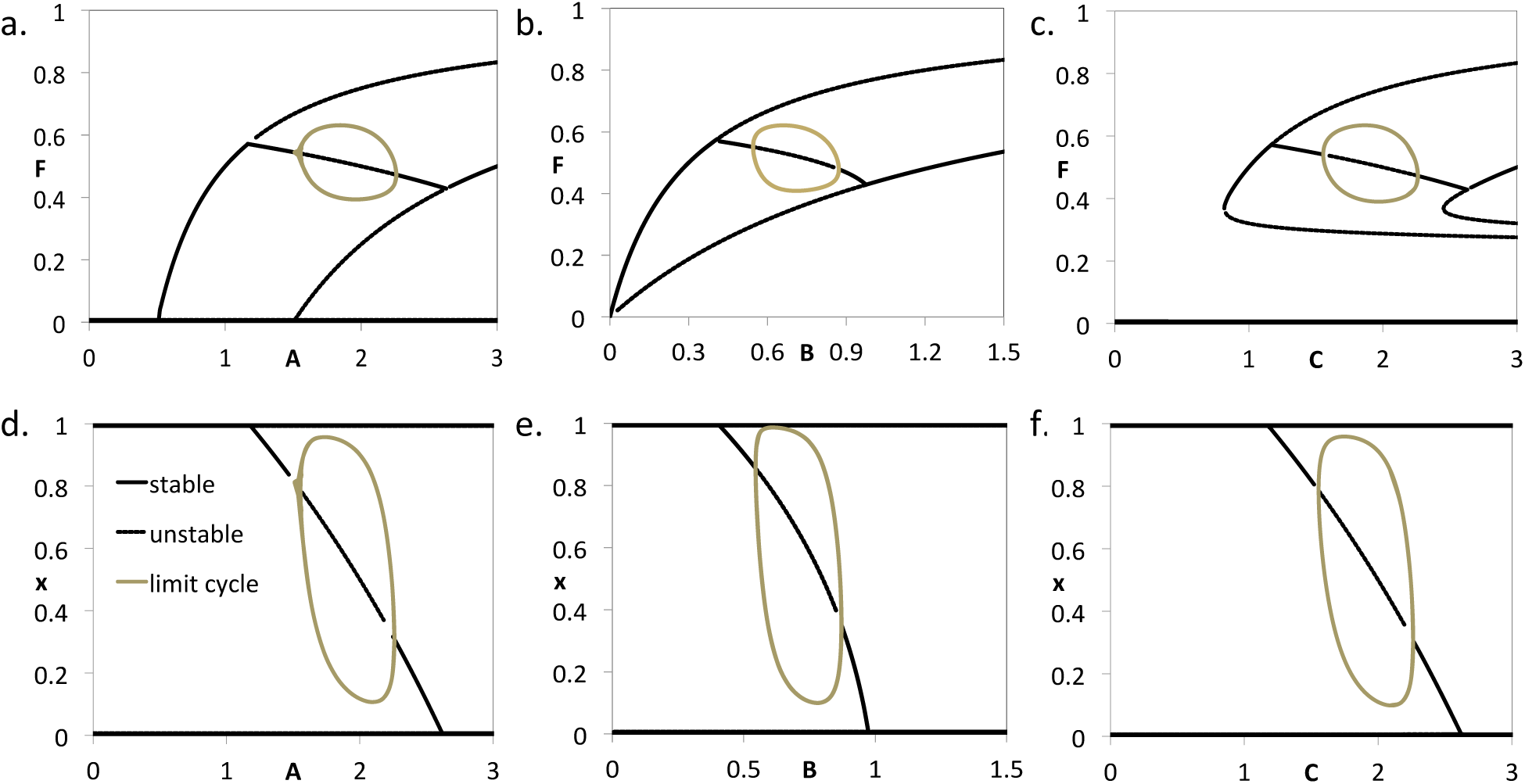
The oscillation in the opinion dynamics at the interior steady state increases largely than in the resource dynamics with large value of the social learning rate. (a, d) Model 1; (b, e) Model 2; (c, f) Model 3. *α* = 5 for Model 1 & 3 and *α* = 8 for Model 2. All other parameters are in Table 2.

## 4. Discussion

Our objective was to understand how different types of natural resource dynamics influence the dynamics of the larger socio-ecological system of which they are a part. We found that the form of natural resource dynamics leaves its imprint in the coupled socio-ecological when human influence is weak. In this regime, social dynamics are dissimilar to ecological dynamics, and ecological dynamics in the socio-ecological system reflect many of the features of ecological dynamics in isolation from human dynamics. We observed how the addition of weak human influence (our baseline scenario) split the interior stable branches into two separate branches, one corresponding to *x* = 0 and one corresponding to *x* = 1, with a small intermediate regime where 0 < *x* < 1. However, the new branches followed the same general trend as in the isolated system, and bifurcation types such as fold bifurcations carried over.

However, the imprint of the natural systems was increasingly obscured as human influence was strengthened. Stronger human influence expanded the size of the parameter regime where the human population has a heterogeneous opinion structure (0 *< x <* 1). In this parameter regime, the ecological dynamics under the three types of natural dynamics diverged from their form in the isolated ecological system, and they began converging to a similar pattern in all three models. In particular, we observed how growing human influence modulated resource levels across the intermediate parameter regime. As a result, increases in parameter values such as the growth rate or the death rate caused a counteractive response by the human population. This causes resources levels to have only a small dependence on growth/removal parameters. However, when social learning is fast enough, all three models exhibit stable limit cycles in natural dynamics. Moreover, social dynamics also follow this pattern of stable limit cycles. Thus, social and ecological dynamics mirror one another when human influence is sufficiently strong. This echoes coupling-induced synchrony observed in many physical systems, such as coupled oscillators. Outside of this intermediate range, we recovered *x* = 0 or *x* = 1, leaving no room for an adaptive human response as parameter values are modified.

In CPR problems, support for conservationism depends not only upon the abundance of the resource but also upon the resource growth rate. Our model showed how a low resource growth rate (or inflow rate, for Model 2) is associated with cooperation whereas a high growth rate is associated with defection. Similarly, a low resource removal rate is associated with defection and a high removal rate with cooperation. This has also been observed in other theoretical models of harvesting in socio-ecological systems Lade et al. (2013). This is driven by the fact that defectors can get more benefit than cooperators as the resource growth rate increases.

Our tentative finding that socio-ecological dynamics converge to a common pattern where social and ecological dynamics strongly mirror one another–irrespective of the type of natural dynamics–has implications for our understanding of socio-ecological dynamics, including how they respond to policy interventions and other external shocks, as well as how we should model them. However, further research is required to explore the robustness of the finding to other relaxations of our simplifying assumptions, and to explore parameter space more systematically. An empirical test of this hypothesis would have to deal with complicated challenges stemming from very different timescales associated with social and ecological systems, and perhaps more importantly, the long-term nature of these dynamics and the presence of other processes that impact the socio-ecological system. In the same vein, future work could also expand our base model by considering the impacts of external policy interventions, adding stochasticity to capture resource extinction events, or exploring the impact of sources of social and ecological heterogeneity such as additional spatial or social structure.

